# Recapitulation of the embryonic transcriptional program in insect pupae

**DOI:** 10.1101/2022.05.15.491985

**Authors:** Alexandra M Ozerova, Mikhail S. Gelfand

**Affiliations:** Skolkovo Institute of Science and Technology; Institute for Information Transmission Problems (Kharkevich Institute), RAS

## Abstract

Metamorphosis of holometabolous insects is predominantly a motionless stage, when no active feeding is observed and the body is enclosed into the pupa. These physiological properties as well as undergoing processes resemble embryogenesis since at the pupal stage organs and systems of the imago are formed. Therefore recapitulation of the embryonic expression program during pupa development could be hypothesized. To assess this hypothesis at the transcriptome level, we have performed a comprehensive analysis of the developmental datasets available in the public domain. Indeed, for most datasets, the pupal gene expression resembles the embryonic rather than the larval pattern, interrupting gradual changes of the transcriptome. Moreover, changes in the transcriptome profile during the pupa-to-imago transition are positively correlated with those at the embryo-to-larvae transition, suggesting that similar expression programs are activated. Gene sets that change their expression level during the larval stage and revert it to the embryonic-like state during the metamorphosis are enriched with genes associated with metabolism and development.

## Background

Hemi- and holometabolous insects differ in the magnitude of physiological and morphological changes during the metamorphosis. In hemimetabolous insects, embryogenesis typically ends up with an adult-like larvae that further develops to the imago through sequential molts causing gradual shifts, with the wings and genitalia appearing during the adult molt. In holometabolous insects, the adult body plan is established at the pupal period, and larval organs and systems are de-differentiated and reorganized during the complete metamorphosis. This is usually accompanied by a more or less radical change in the habitat and feeding strategy. Larvae and adults of the same species do not share food resources, allowing the separation of growth and reproduction in time and space (1).

Metamorphosis is predicted to originate approximately 400 million years ago in early Devonian, when Pterygota has emerged, the insect flight has been invented (2) and complete metamorphosis has evolved to support the ability to fly. Sequential molts require the whole body including the wings to be covered with the cuticle. It makes wings too heavy and almost no extant winged insects undergo molting during the imago stage, an exception being the short-living subimago of the mayfly that undergoes a full molting cycle to become the imago (3).

The larval stage of holometabolous insects has appeared as the arrest of the embryonic program (4) that continues after the pupation. In particular, cells with latent embryonic potential, initially forming so-called imaginal primordia, replace larval cells to form adult organs. The imaginal cells contribute little to the functioning of the larval organism and preserve pluripotency similar to stem cells (5). For example, in *Papilio xuthus* (Lepidoptera), a sophisticated orchestra of transcription factors that regulate the expression patterns of opsins, manifest only after the pupation to build the compound eye (6). On the other hand, some organs undergo dedifferentiation followed by redifferentiation to the adult state. For example, in *Drosophila* (Diptera), syncytial alary muscles de-differentiate to mononuclear myoblasts prior to formation of the adult tissue (7).

Differentiation of stem cells into mature tissues could reuse molecular mechanisms that drive the embryonic development since the gain of new features is based on the upgrade of the existing ones (8). Therefore it could be hypothesized that the pupal gene expression program should resemble the embryonic one due to both differentiation d*e novo* and redifferentiation. Study on the midge *Polypedilum vanderplanki* showed inversion of the transcriptional profile back to the embryonic stage during pupal development (9). Here, we report a comprehensive analysis of the insect developmental transcriptome datasets available in the public domain, aimed to check gene expression similarities between pupae and embryos.

## Methods

### Datasets

Developmental transcriptomic datasets with at least one sample originating from each of the four major stages (embryonic, larval, pupal, adult) in the holometabolous insect development were analyzed. The collection includes ten species from four orders (Diptera, Hymenoptera, Lepidoptera, Coleoptera). For *Drosophila melanogaster*, both RNA-seq and microarray datasets are available, and RNA-seq only for all other species, see Table 1 and Supplementary Tables 1-10 for details.

**Table 1.**
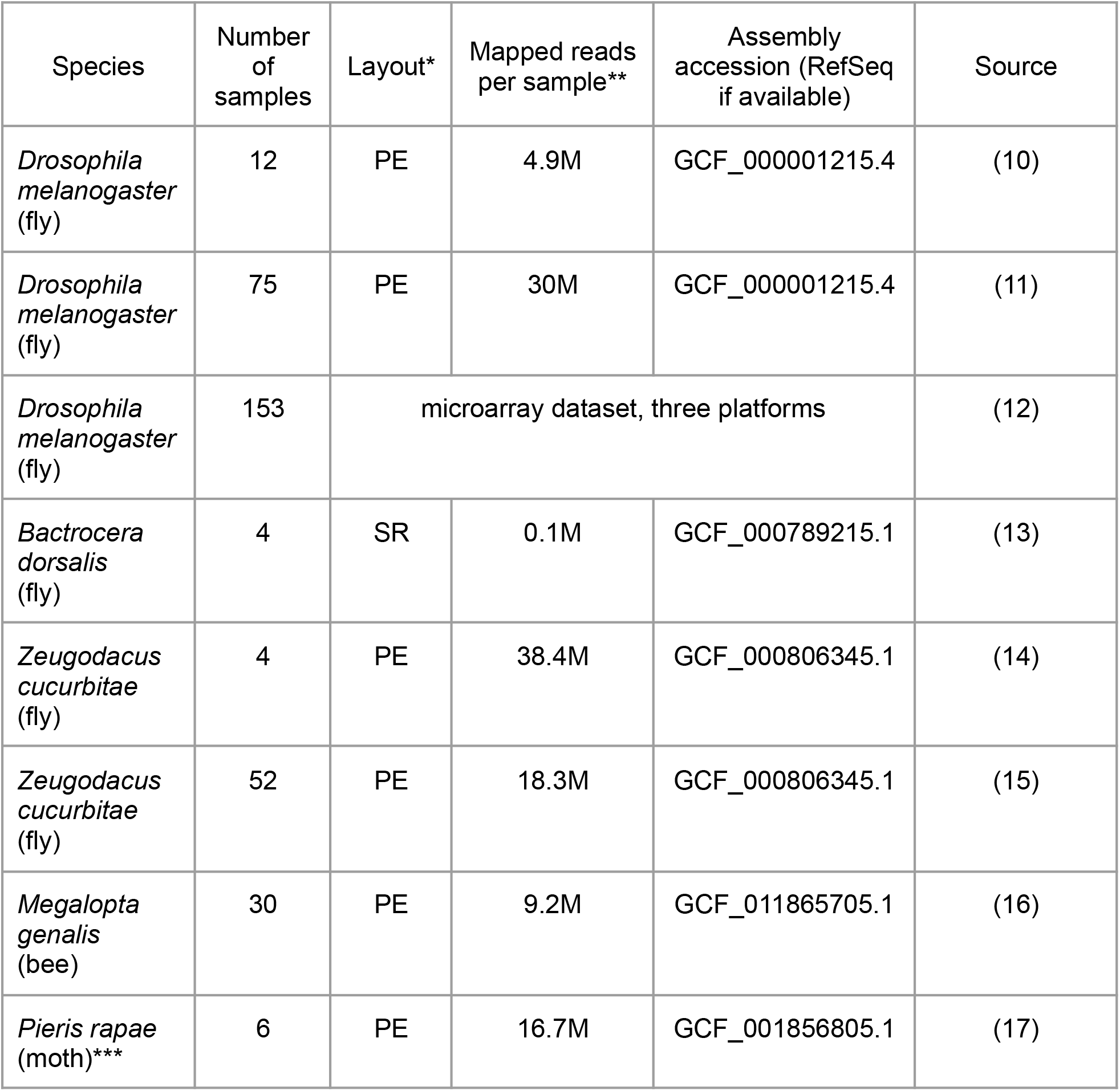
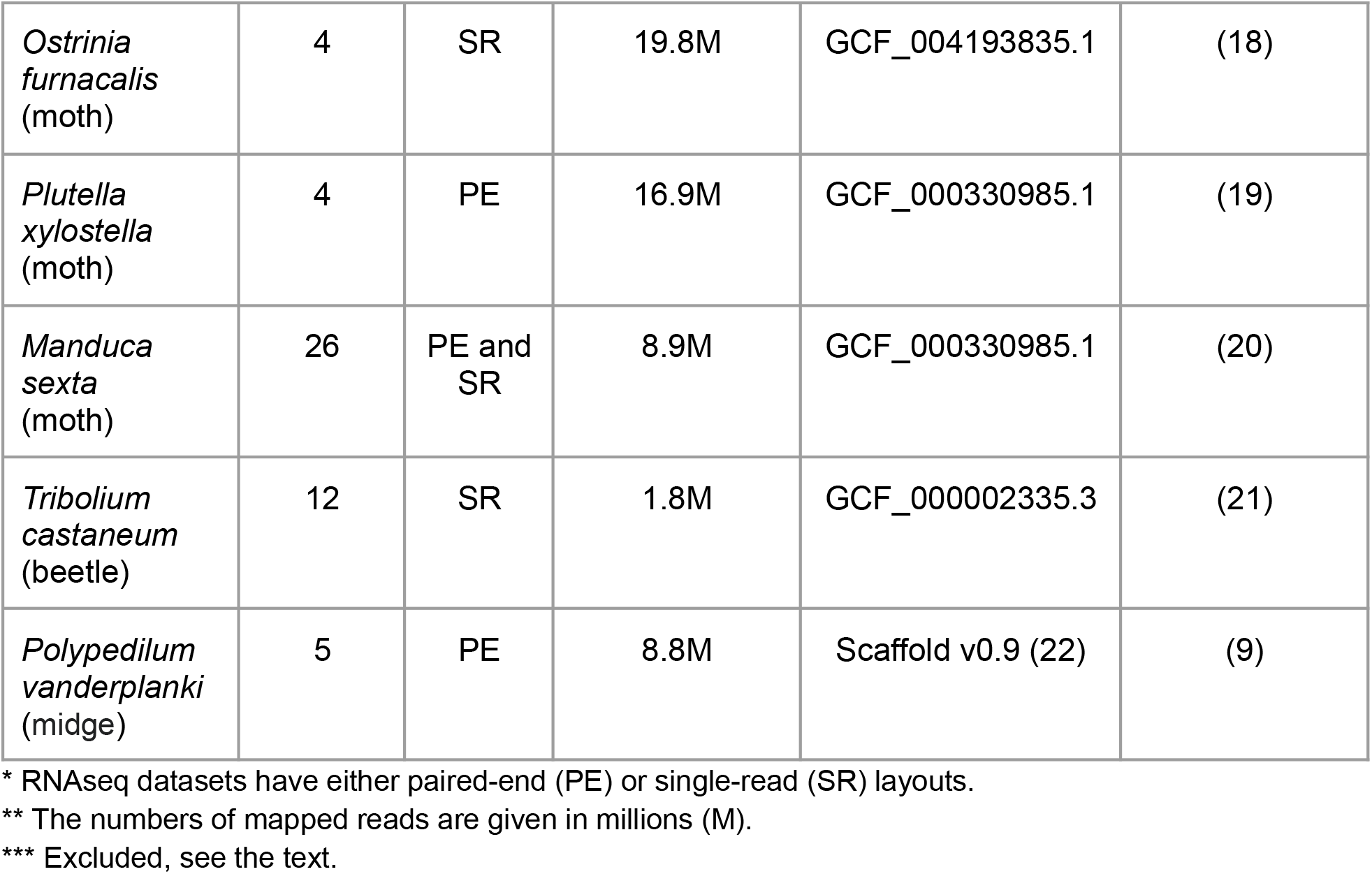
Datasets.

Mature females contain eggs, therefore full-body transcriptomes have a strong signal from the eggs, yielding a high correlation of the female samples with the embryonic state. To avoid these confounding effects, the female samples were excluded from the analysis.

After the initial analysis, the *Pieris rapae* dataset was excluded due to insufficient data for the embryonic sample, the latter being an outlier with only 0.2M uniquely mapped reads, compared to about 13M reads for each other sample.

The *Tribolium castaneum* dataset comprises three replicates for each developmental stage. Two of the pupal samples had exactly the same set of raw reads, therefore one copy was excluded.

### RNAseq preprocessing

RNA sequencing reads were downloaded from the NCBI sequence read archive in the fastq format. Low-quality reads were eliminated, adapters and low quality nucleotides were trimmed using the fastp tool (23). Reads were mapped to the reference transcriptomes (listed in Table 1 for each species) using the Kallisto tool (24). The FPKM (Fragments Per Kilobase Million) normalization was used for the downstream analysis.

### Microarray dataset

Processed data containing log-transformed ratios between channels were downloaded from the NCBI GEO database (accession GSE3286) for three platforms separately.

### GO terms annotation

*Drosophila melanogaster* gene ontology (GO) annotation was downloaded from the AmiGO 2 database (25).

InterProScan with the default parameters was used to predict GO terms for other species (26). Protein sequences from the respective assemblies (see Table 1) were used.

Genes were assumed to be related to development, if the Gene Ontology (GO) term “developmental process” (GO:0032502) or its descendant terms were predicted to be associated with the gene. Genes predicted to have the GO term “metabolic process” (GO:0008152) or its descendants were regarded as metabolism-related.

### Across-stages similarity

Similarity between stages was measured by the Spearman correlation coefficient of the FPKM values.

### Random sampling

To assess the influence of particular gene subsets on the observed transcriptome changes, random sampling was performed. At that, gene sets of the same size were sampled, and for each sample the correlation coefficient was computed. The quantile of the tested gene set was estimated from the obtained distribution.

### Gene profile clustering

For datasets with data available for four main stages only, there are 27 possible patterns of gene expression (it could increase, decrease or remain the same during each of three transitions between stages). A gene was assigned to the pattern with which it has the highest correlation.

The transcriptome datasets with more than four time points were hierarchically clustered with the Spearman correlation coefficient as the distance metric. Hierarchy algorithm from the *scipy* package was used (27).

### Gene ontology term enrichment analysis

Python package *goatools* was used to identify significantly enriched GO terms (28) using the default parameters with an adjusted p-value threshold equal to 0.05.

### Visualization

Python 3.7, *matplotlib* and *seaborn* packages were used for the visualization. Plots for the GO enrichment results were generated using the REVIGO tool (29) and the R *ggplot* package.

## Results and Discussion

### Intra-species comparison

Intra-species comparisons across developmental stages allowed us to compare the transcriptome profiles at several distinct time points. A monotonic development results in gene expression patterns at each particular stage being closest to the immediately previous and following developmental stages, yielding a decrease of the similarity with the increase of the time interval between the time points. Indeed, this behavior is observed for the embryonic and larval stages. The correlation coefficient decreases for relatively more distant stages as seen in the pairwise correlation heatmap for the detailed *Drosophila melanogaster* dataset (Figure 1). In that case, high Spearman correlation coefficients are concentrated near the diagonal.

**Figure 1.**
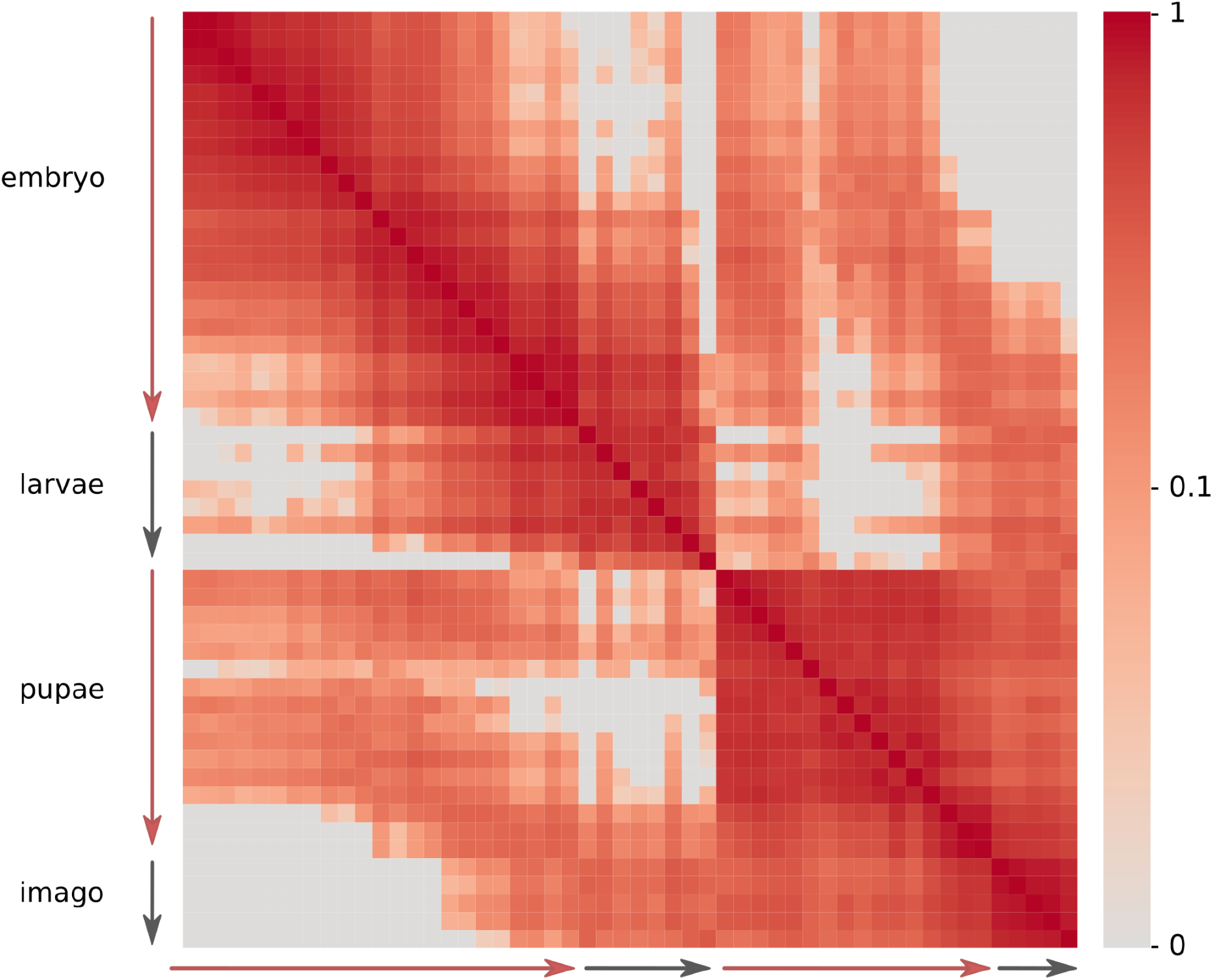
Pairwise correlation coefficient heatmap for *Drosophila melanogaster* developmental stages. The expression data are from (12).

However, this monotonic development is interrupted during the pupation suggesting drastic changes in the transcriptome profile. Gene expression levels at early pupal stages are closer to the embryonic profiles rather than to the larval ones. The monotonic development is restarted at some point during the pupal development, extending to the adult stages, so that high correlation coefficients are again observed close to the diagonal of the matrix.

Datasets from several other insect species, though less detailed, demonstrate the same overall pattern with pupal transcriptomes being more similar to the embryonic ones than to larval or adult ones. Sample heatmaps for moth *Ostrinia furnacalis* and beetle *Tribolium castaneum* are presented in Figure 2 (left). Heatmaps for other datasets see in Supplementary Figure 1.

**Figure 2.**
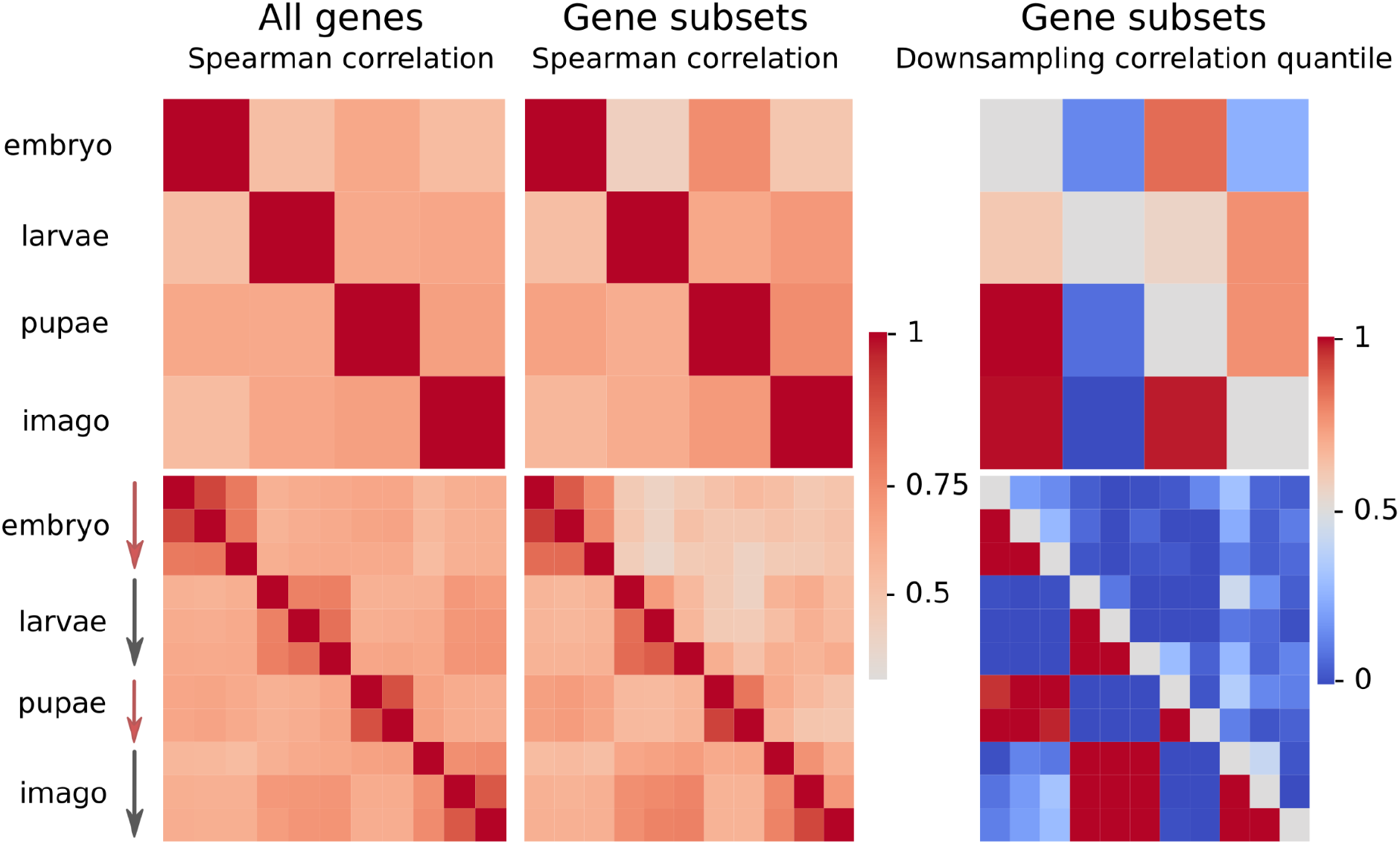
Pairwise correlation analysis for *Ostrinia furnacalis* (top) and *Tribolium castaneum* (bottom). The Spearman correlation coefficients, calculated considering all genes (left), development-associated gene (middle, the upper triangle), metabolism-associated (middle, the lower triangle) and results of the random sampling analysis (see Methods) considering the development-associated gene subset (right, the upper triangle) and the metabolism-associated gene subset (right, the lower triangle)

The effect of increased similarity between embryo and pupa compared to the embryo-larvae similarity is observed in several more datasets (Figure 3a, left), therefore it is not restricted to the *Drosophila* genus or the Diptera order. However, for some species pupae do not resemble embryos, an example being *Manduca sexta* (Figure 3, blue line). This could be explained by the fact that the *M. sexta* samples were collected from the whole body for early developmental stages and from several specific tissues for later stages. This makes direct transcriptome comparisons less reliable, since differences could be tissue-specific regardless of the developmental stage. In other cases, a possible explanation is that early or late pupae have been collected, closer to the adjacent larval or adult stages, respectively.

**Figure 3.**
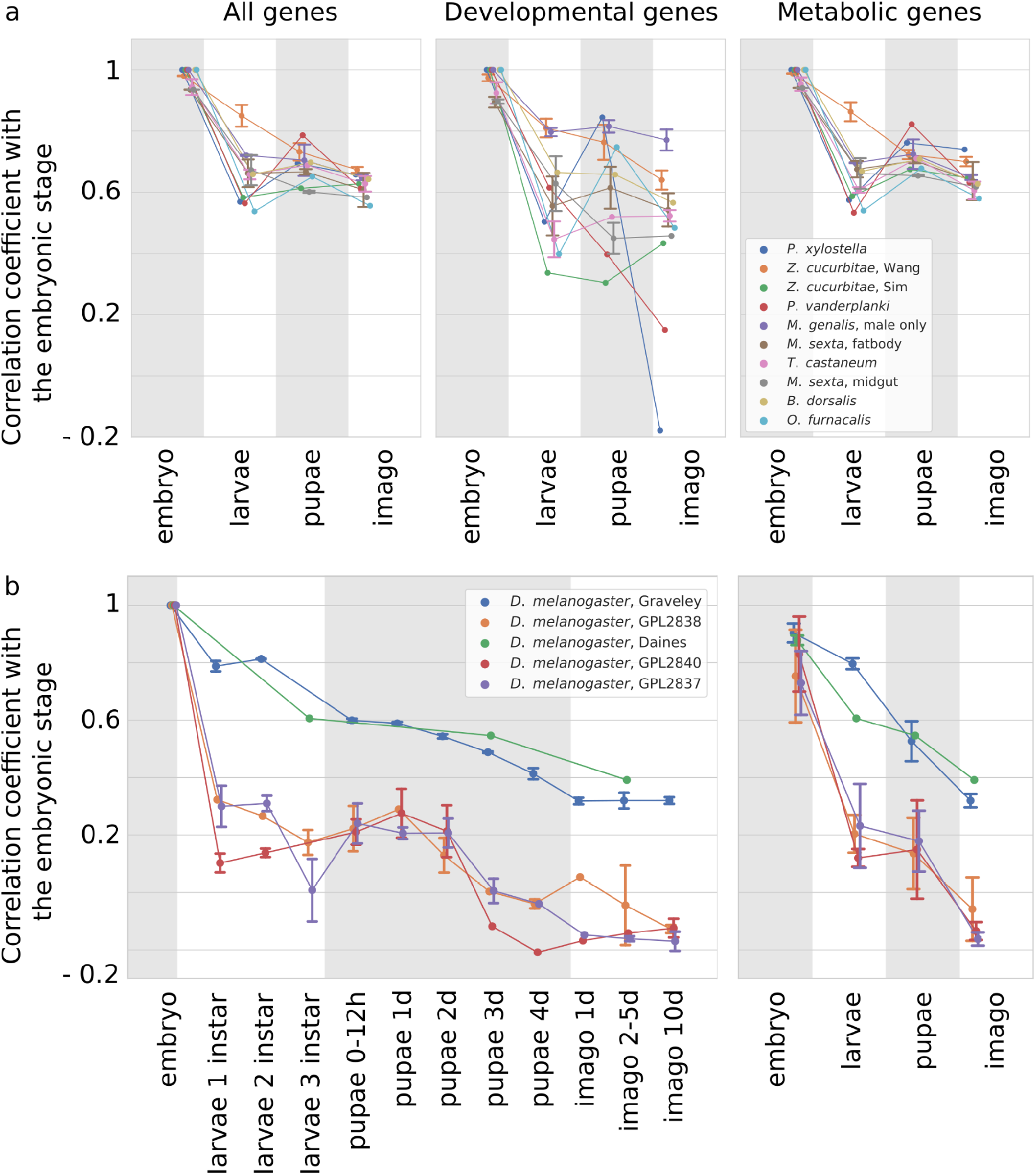
Correlation between the developmental stage transcriptional profiles (horizontal axis) and the embryonic profile. (a) All species excluding *Drosophila melanogaster*. (b) *Drosophila melanogaster* datasets. Quartiles are shown for datasets with more than one sample collected during each of the major stages

The latter explanation is supported by three *D. melanogaster* microarray datasets (Figure 3b). Indeed, while middle pupal stages are more similar to the embryonic stages, late pupae are more similar to adults.

*D. melanogaster* datasets fall into two groups depending on the technology used to generate the source data. The RNA sequencing datasets (orange and purple lines in Figure 3b) demonstrate monotonic decrease of similarity with the embryo in the course of development.

### Functional subsets of genes

To understand the molecular basis of the observed pattern, functional subsets of genes were considered. During pupal development, tissues are reorganized or even developed *de novo* from stem cells like in embryogenesis. From the lifestyle point of view, the pupa also resembles the embryo since it is motionless and lacks active feeding. Therefore, genes with Gene Ontology (GO) terms related to “developmental process” (GO:0032502) and “metabolic process” (GO:0008152) were tested to account for the observed effect.

For *D. melanogaster* RNA-seq datasets, the analysis of subsets yields largely similar results. For two detailed microarray datasets, genes associated with metabolic processes demonstrate higher correlation between the embryonic and pupal samples, however the range of the interval between quartiles makes the observation unreliable. Similarity between embryo and pupa is higher, when calculated based on gene subsets, rather than the entire dataset for the *O. furnacalis* (Figure 2a, middle). To test the statistical significance of the finding, random sampling was performed (see Methods). A high quantile of the observed correlation coefficient is expected for gene subsets strongly influencing the effect. On the contrary, gene subsets that have expression patterns following the average trend would have quantile values close to one half. High correlation between embryonic and pupal transcriptomes in *O. furnacalis* data is supported by quantile values (Figure 2, right top), suggesting that both development and metabolism-associated genes are collinearly expressed during these stages.

On the other hand, in *T. castaneum*, the expression of development-associated genes does not follow this trend (Figure 2b, middle). At that, it should be noted that *D. melanogaster* is the only analyzed species with verified GO terms annotation from a dedicated database, while the GO annotation for other species is predicted using InterProScan. Moreover, only 1% of all proteins are annotated as associated with GO:0032502 (“developmental process”), leading to noisy results (Figure 3, middle top). On the contrary, as many as 40% of proteins in each species are assigned with GO:0008152 (“metabolic process”) and hence the correlation coefficients naturally are close to those obtained using the whole datasets (Figure 3, right top). However, for both types of subsets there are species with an enhanced effect.

### The inversion of the transcriptome pattern

As described above, the transcriptome profile of holometabolous insects tends to revert to the embryonic state during metamorphosis. This can be seen not only from direct comparison of transcriptomes on several developmental stages, but from the analysis of transcriptome changes during transitions between adjacent stages. Indeed, in some cases changes in gene expression that happen at the pupa-to-imago transition recapitulate the egg-to-larva transition. For example, the left part of Figure 4 shows changes in gene expression for the *O. furnacalis* and *T. castaneum* datasets. Positive correlation between fold-change values is observed in 75% of the datasets. 40% of them have correlation coefficient higher than 0.1, suggesting that the pupal developmental program that drives (re-)formation of tissues and organs indeed dynamically recapitulates the embryo differentiation.

**Figure 4.**
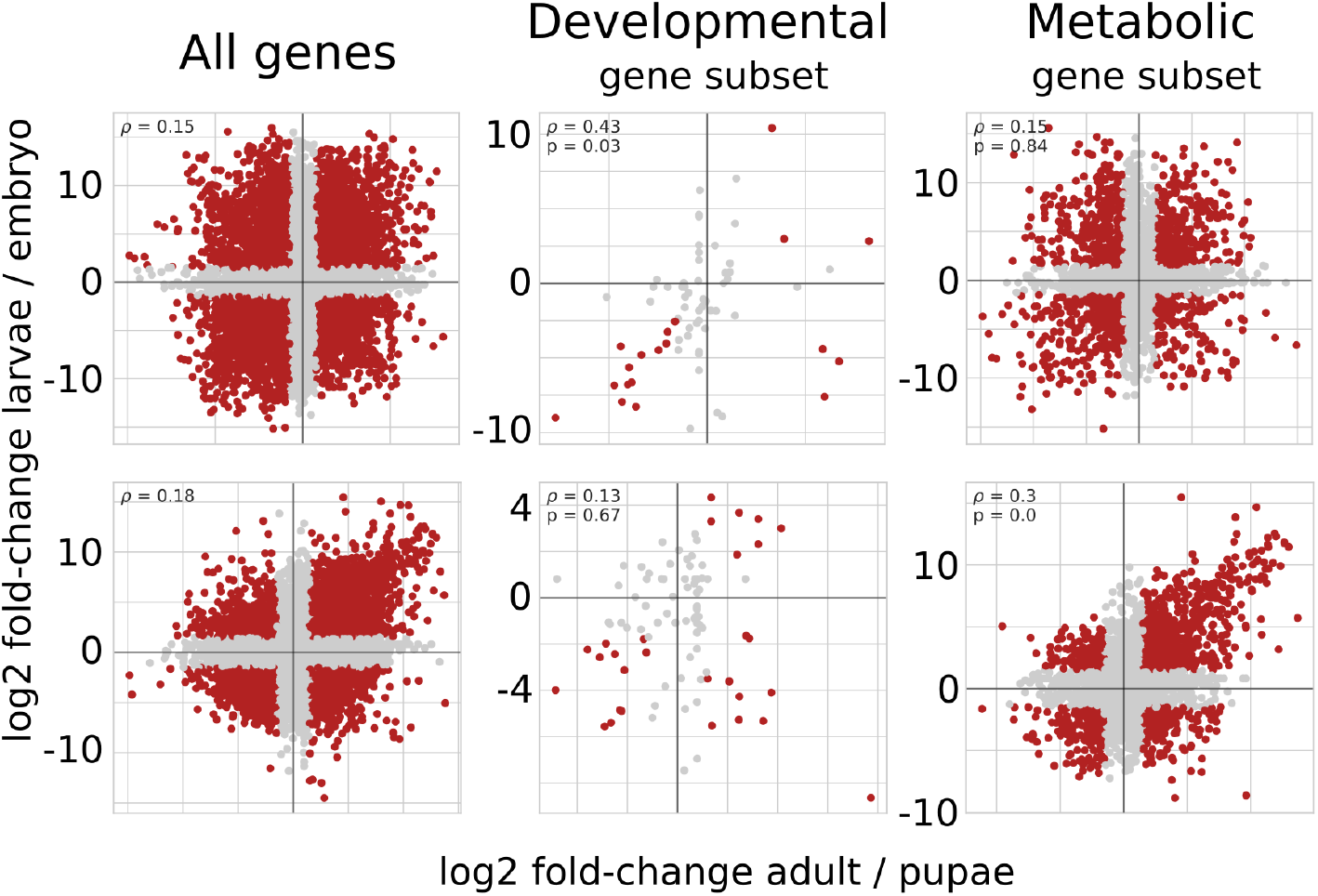
Changes in transcriptome profiles during the transition from the pupal to the imago stages compared to the transition from the embryonic to the larval stages for *Ostrinia furnacalis* (top) and *Tribolium castaneum* (bottom). Each dot represents one gene, with the embryo-to-larvae expression fold-change on the y-axis and the pupa-to-imago fold-change on the x-axis (log scales). Three gene sets are considered: all genes (left), development-associated genes (middle), metabolism-associated genes (right). Genes with significant LCF (greater than 1.5) are shown in red.

The effect of synchronized changes in expression patterns during embryo and pupal eclosion could be explained by monotonous processes occurring in the complete course of the development. However it could not be the main explanation, since in pairwise comparisons gradual changes are not observed and the pupal transcriptome is more similar to the embryonic rather than larval one.

For *O. furnacalis*, the development subset demonstrates higher correlation between fold-changes (middle top of the Figure 4). Downsampling p-value supports this observation since only 1% of random gene subsets show higher correlation. However, due to the low number of genes with predicted link to development, a significant effect is seen in only five datasets with positive correlation (Figure 5, left).

**Figure 5.**
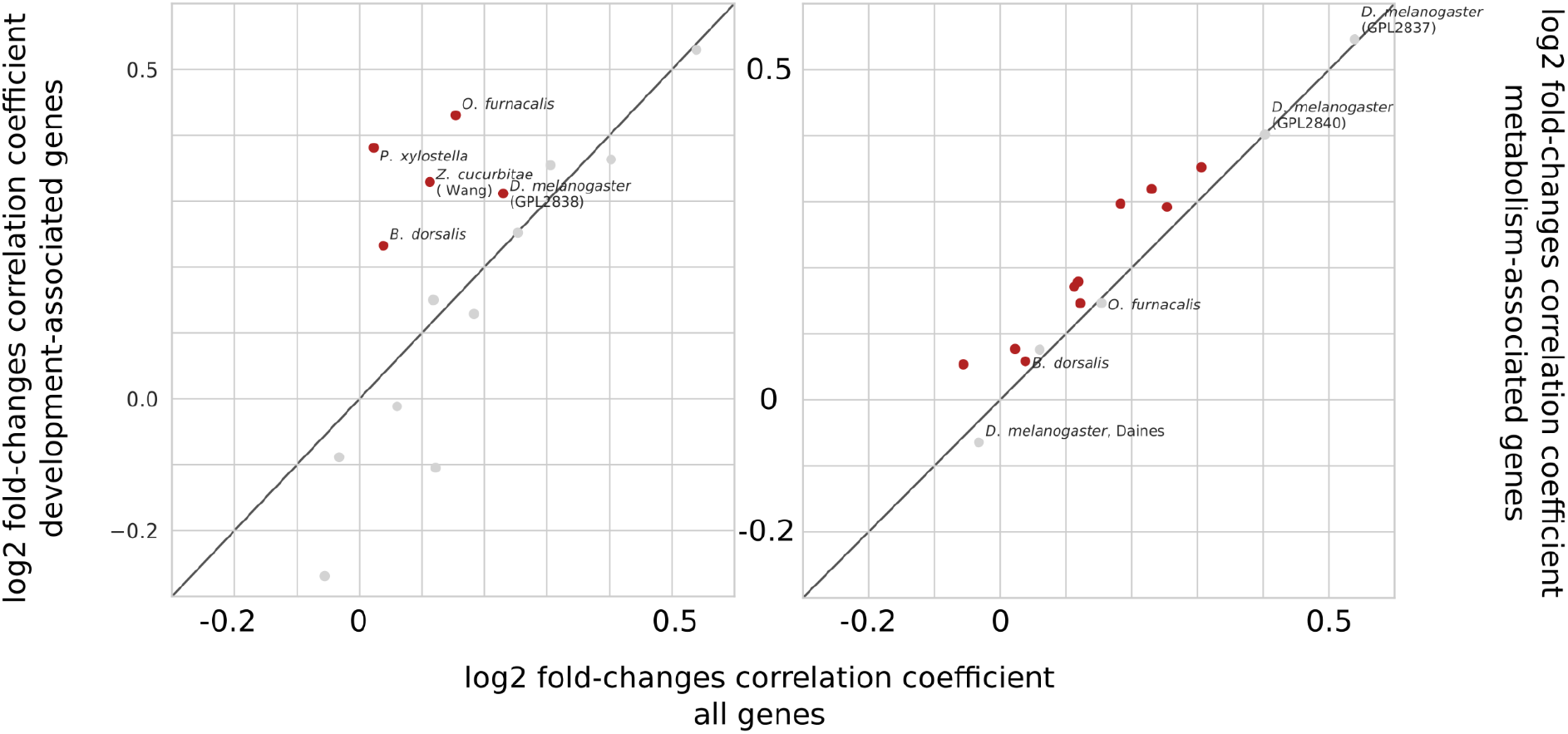
Correlation between transcriptome transitions from the embryo to the larva and from the pupa to the adult for all species. The correlations are calculated on the entire dataset (horizontal axis), the development-associated subset (according to gene ontology, vertical axis, left) and the metabolism-associated subset (vertical axis, right). Dots corresponding to the datasets with the statistically significant correlation coefficient are shown in red.

Higher than average fold-change correlation for metabolism-associated genes is observed across more datasets (Figure 5, right), and it is statistically significant for most of the species. Therefore metabolic genes could partially drive the recapitulation.

Genes that drive recapitulation should have a zig-zag-like pattern of gene expression during development. To identify such genes, clustering of the expression profiles was performed. Datasets with one time point measured for each of the main stages (embryo, larva, pupa and imago) are scored by a correlation coefficient with one of the possible 27 artificial trajectories (with up/same/down steps).

More detailed datasets were clustered hierarchically (see Methods). The *Manduca mexta* dataset was not considered at this step since 72% of its samples correspond to the larval stage and therefore the cluster diversity is dominated by gene expression changes during the larval development. For other datasets, clusters with the pattern of similar expression in the embryonic and pupal samples (zig-zag-like pattern) were selected for further analysis (Supplementary Figure 2). The set of genes with expression that increases while entering the larval stage and then decreases after the pupation is enriched with several classes of metabolism and development-associated terms (Figure 6).

**Figure 6.**
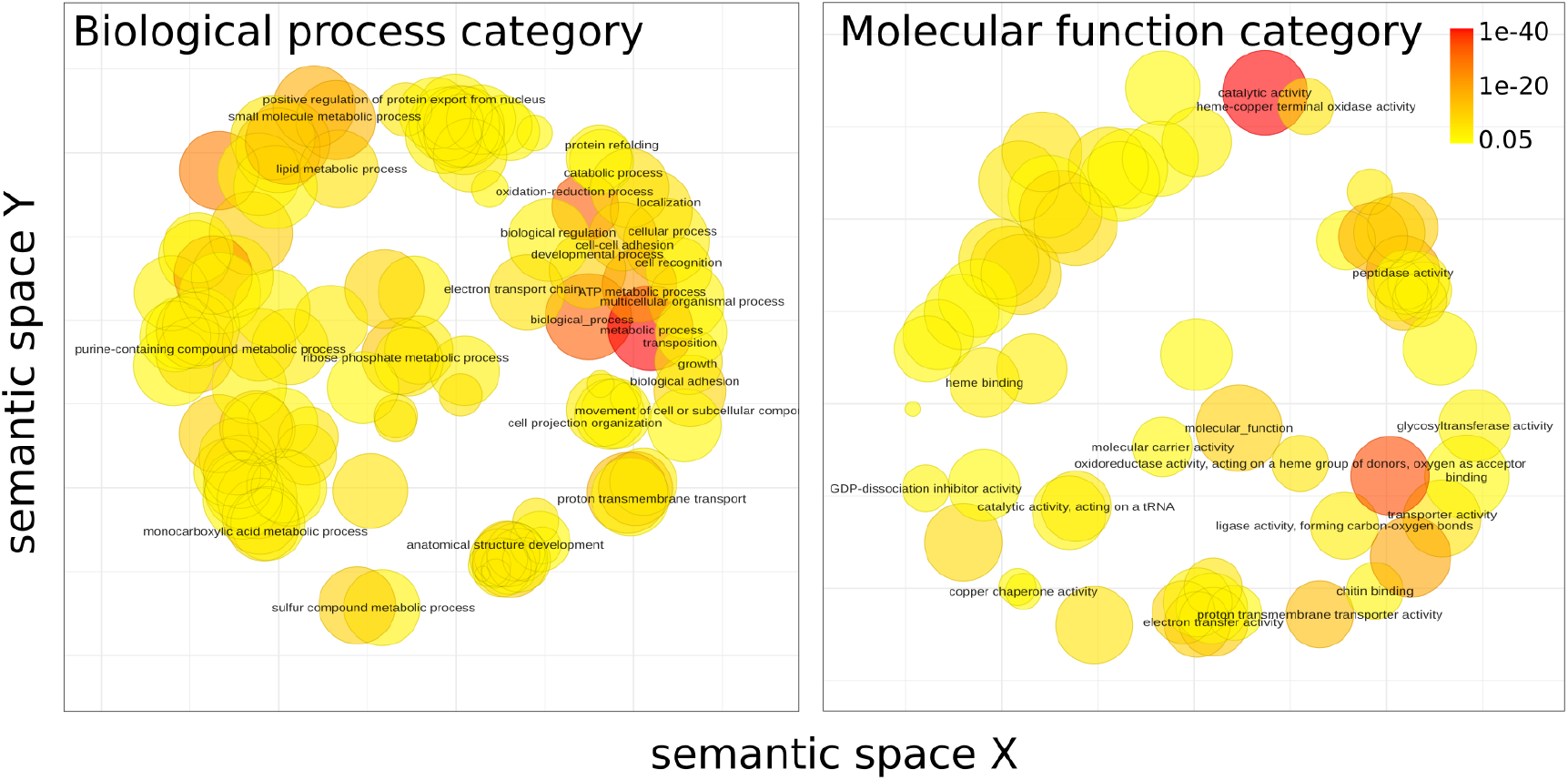
Gene ontology enrichment for genes primarily expressed during the embryonic and pupal stages. GO terms are projected so that semantically close terms are spatially close. The color represents adjusted p-value, multiplied over all the datasets. The size of circles reflects the log-transformed number of the term in the EBI GOA database (29).

Terms similar to *purine containing compound metabolic process* (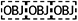GO:0072521), *ribose phosphate metabolic process* (GO:0019693), *ATP metabolic process* (GO:0046034), *electron transport chain* (GO:0022900), *proton transmembrane transport* (GO:1902600) and *oxidation-reduction process* (GO:0055114) in the semantic space for biological processes category suggest a high rate of energy-generation and consumption during both embryogenesis (30) and pupa maturation to build organs and tissues.

The enriched *peptidase activity* (GO:0008233) molecular function could be involved in several processes. For example, matrix metalloproteinases regulate trachea and intestines development of embryo and pupal morphogenesis of *Tribolium castaneum* beetle (31) and caspases along with other proteases are key players in the induced cell death. During metamorphosis, it is essential for remodeling of the larval tissue (32). Peptidases also balance proliferation, being, e.g., crucial players in development of the tracheal system (33) or neuroblasts (34) in *Drosophila melanogaster*.

*Cell-cell adhesion* (GO:0098609), *multicellular organismal process* (GO:0032501) and *anatomical structure development* (GO:0048856) terms are frequent among active genes during the embryo and pupal stages. In particular, these terms were enriched in the respective cluster in the *D. melanogaster* RNA-seq dataset (11) (Figures 7a and 7b, respectively). The correlation heatmap for this cluster features a distinct diagonal suggesting each stage is similar to the ones that are close in time (Figure 7c). However, there is a prominent diagonal in the embryo-pupae submatrix suggesting involvement of similar processes.

**Figure 7.**
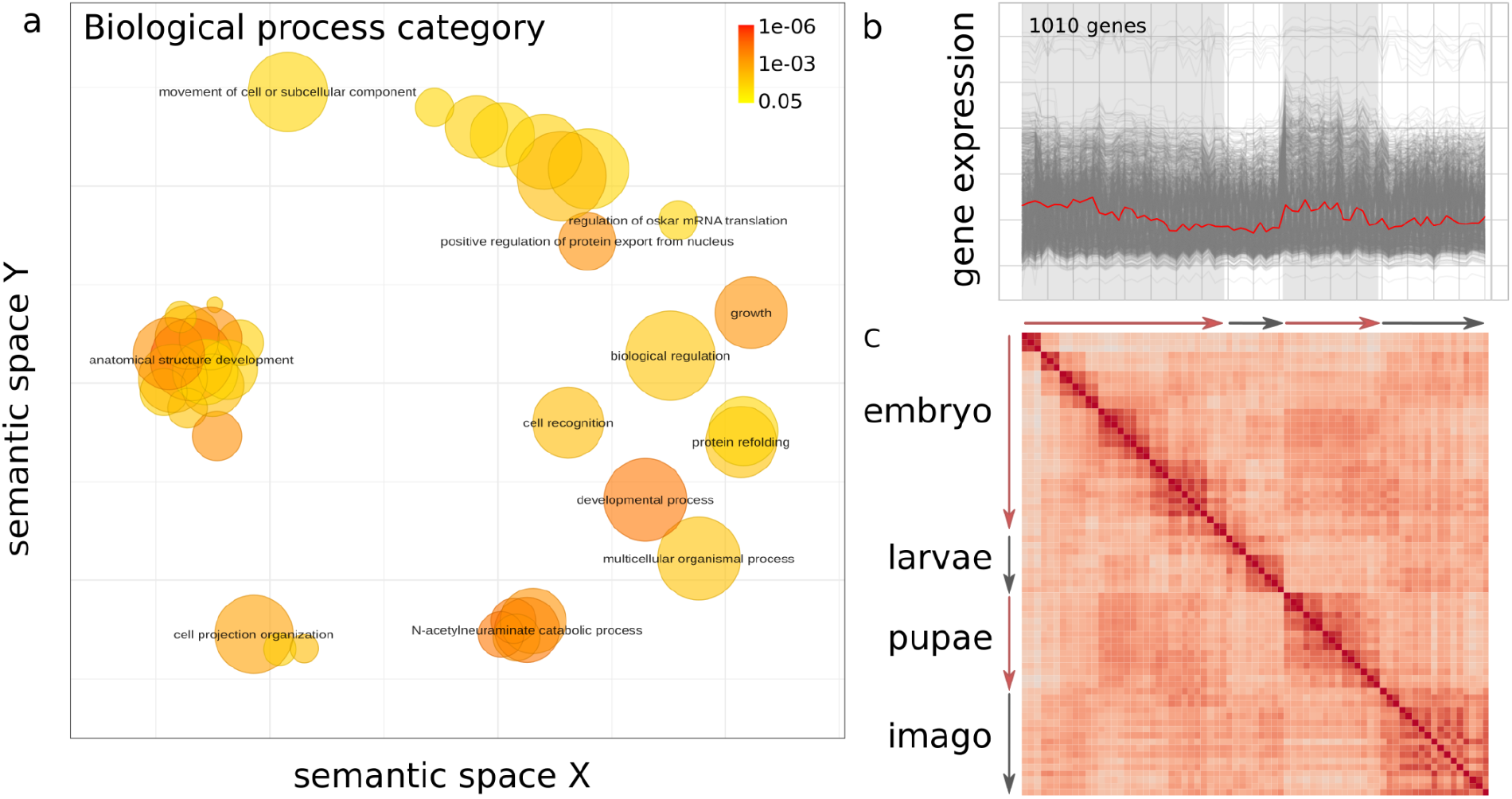
Analysis of the zig-zag cluster in the *Drosophila melanogaster* dataset. (a) Gene ontology enrichment results for the genes comprising the cluster. (b) Gene expression patterns across the development for the zig-zag cluster. (c) Pairwise Spearman correlation coefficients for genes from the zig-zag cluster.

## Conclusions

Several datasets indeed show an increased similarity between the embryonic and pupal stages on the gene expression level when compared with the embryo-larvae transcriptome pairs. Sets of genes changing their expression level during the larval stage and returning to the embryonic state during the metamorphosis are enriched with genes related to energy metabolism and multicellular organism development.

Gene expression changes at transition from the embryonic to the larval stage for some datasets are correlated with changes between pupa and imago, suggesting similarity of transcriptional programs during embryonic development and pupal maturation. Separate analysis of metabolism-associated genes and genes related to the development enhances the observed effect for most datasets.

However, some datasets do not follow the pattern of embryonic expression recapitulation during morphogenesis. This might be due to the timing of collected pupal stages, as early pupae naturally resemble late larvae, while late pupae are similar to imagos. Still we consider the hypothesis to be tentatively confirmed and submit it for detailed experimental validation.

## Acknowledgements

This study was supported by grant 20-54-81007 from the Russian Foundation for Basic Research.

We are grateful to Georgii Bazykin, Ekaterina Khrameeva and Mikhail Moldovan for useful discussions.

