## Supplementary Figure 1 for "Recapitulation of the embryonic transcriptional program in insect pupae"

### Supplementary Figure 1. Pairwise correlation analysis.

The Spearman correlation coefficients, calculated considering all genes (left), development-associated gene (according to gene ontology; middle, the upper triangle), metabolism-associated (middle, the lower triangle) and quantiles in downsampling analysis considering the development-associated gene subset (right, the upper triangle) and the metabolism-associated gene subset (right, the lower triangle)

*Drosophila melanogaster*, Daines

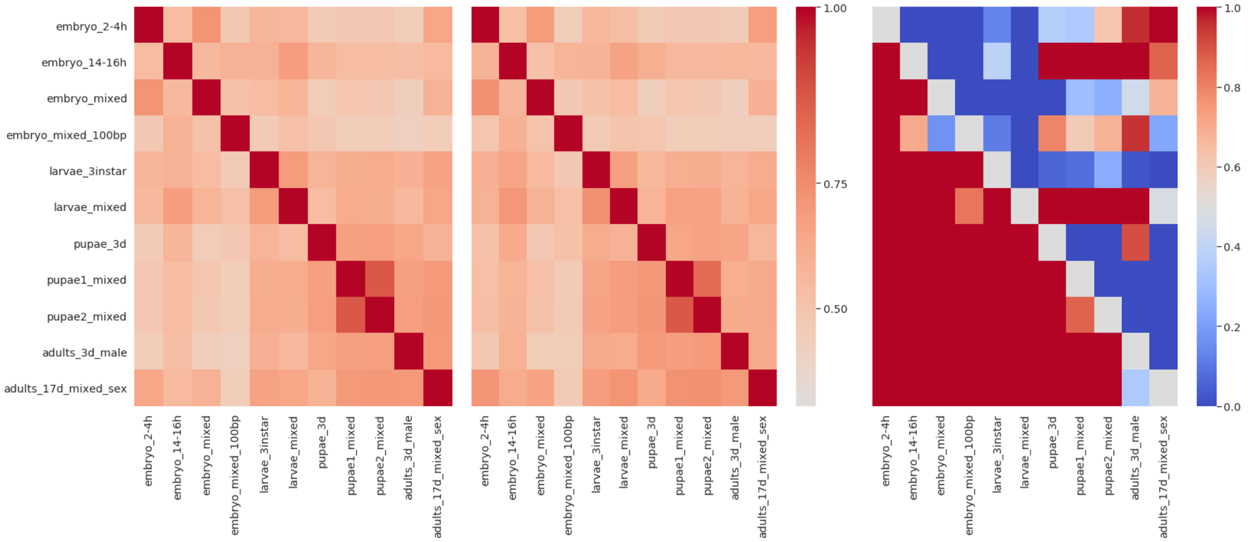


*Drosophila melanogaster*, Graveley

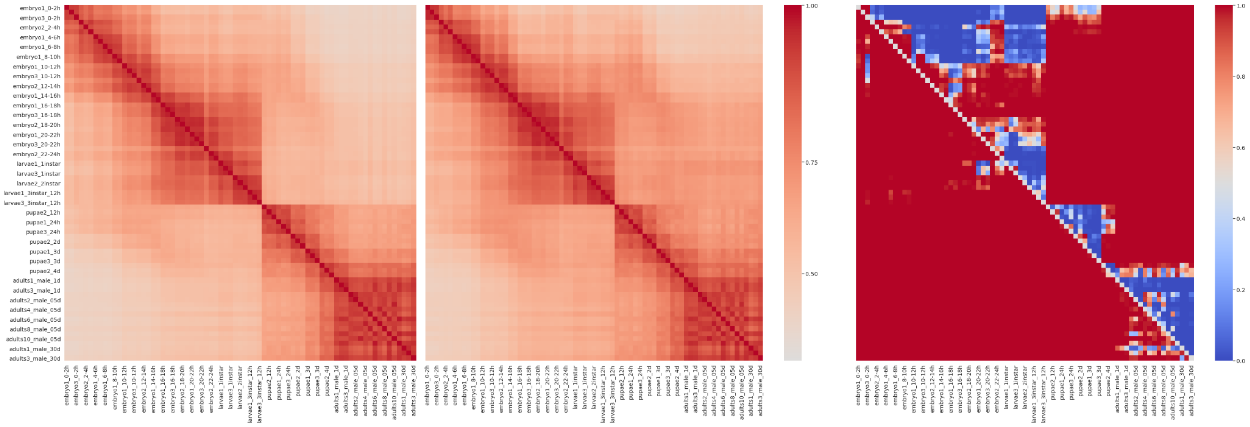

#### *Drosophila melanogaster*, Arbeitman, GPL2837

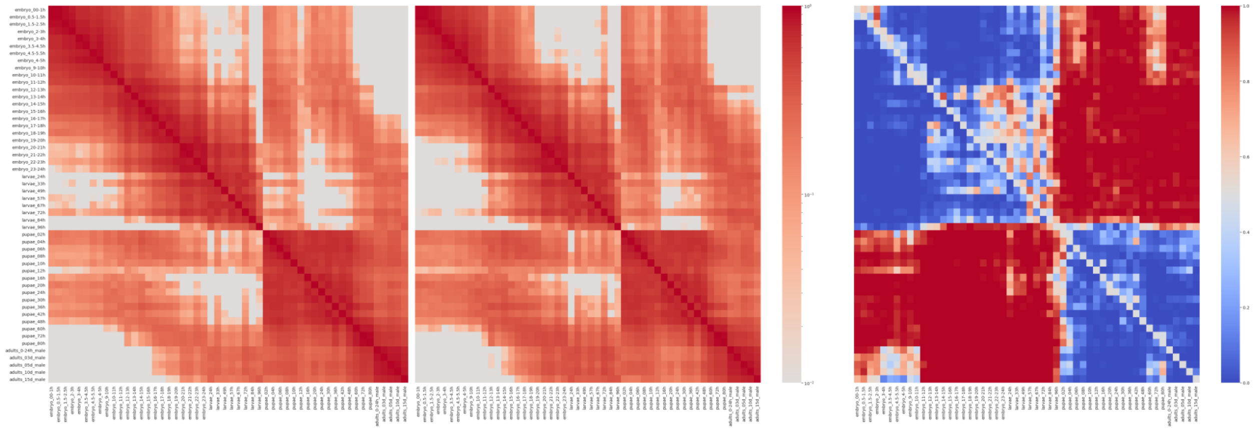

#### *Drosophila melanogaster*, Arbeitman, GPL2838

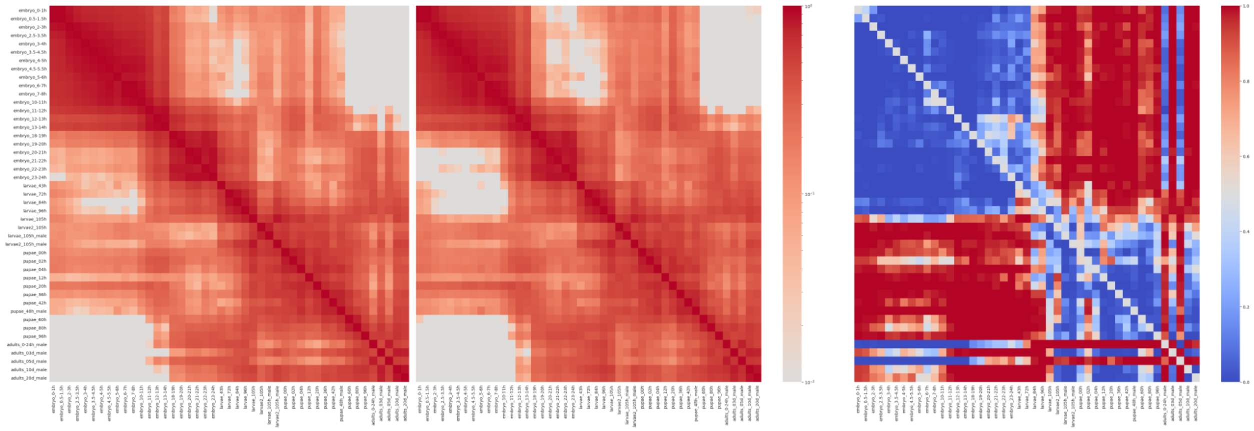

#### *Drosophila melanogaster*, Arbeitman, GPL2840

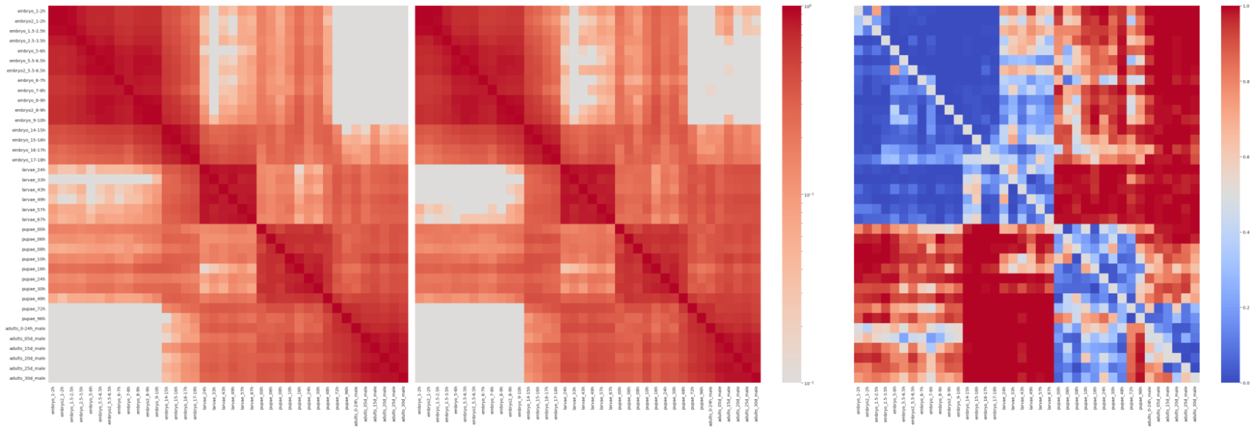

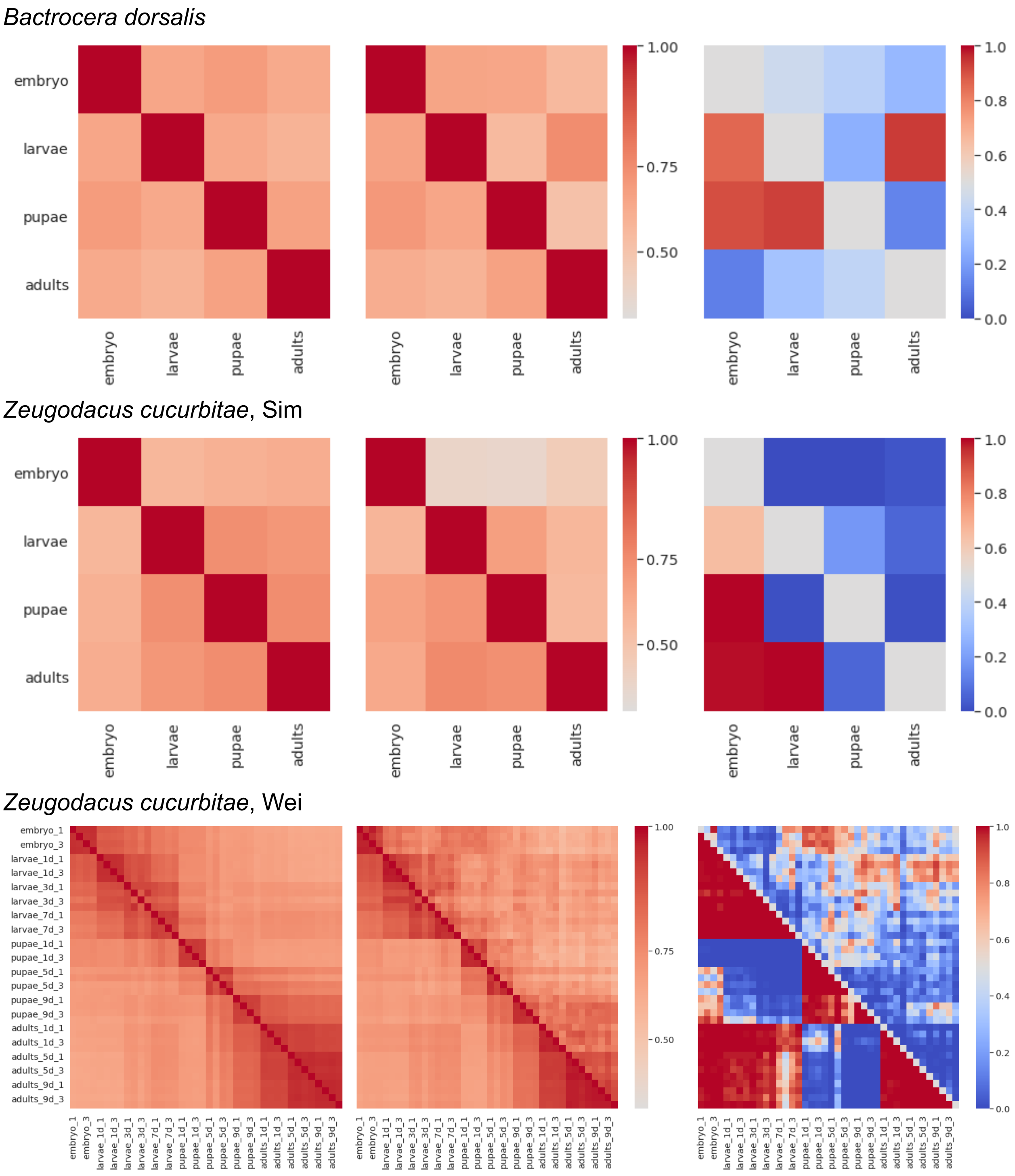
Bactrocera dorsalis
Zeugodacus cucurbitae, Sim
Zeugodacus cucurbitae, Wei
embryo
larvae
pupae
adults
1.00
0.75
0.50
1.0
0.8
0.6
0.4
0.2
0.0
embryo_1
embryo_3
larvae_1d_1
larvae_1d_3
larvae_3d_1
larvae_3d_3
larvae_7d_1
larvae_7d_3
pupae_1d_1
pupae_1d_3
pupae_5d_1
pupae_5d_3
pupae_9d_1
pupae_9d_3
adults_1d_1
adults_1d_3
adults_5d_1
adults_5d_3
adults_9d_1
adults_9d_3

*Megalopta genalis*

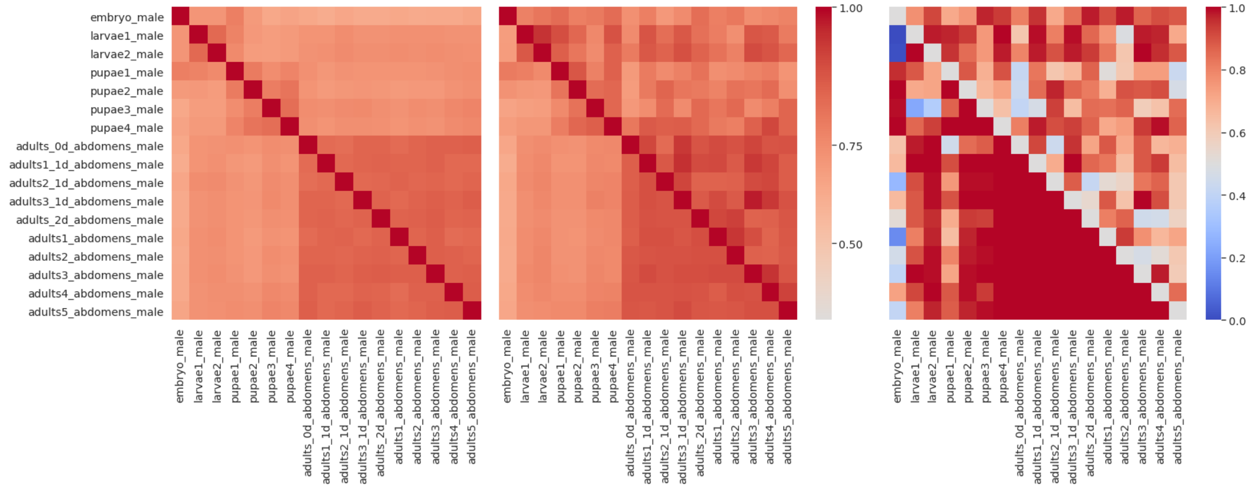


*Plutella xylostella*

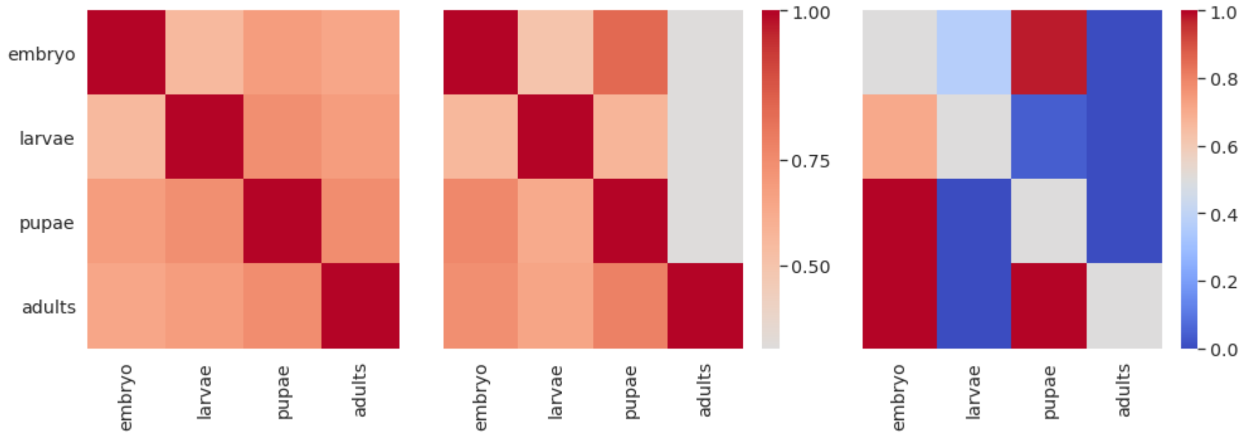


*Manduca sexta*, midgut samples

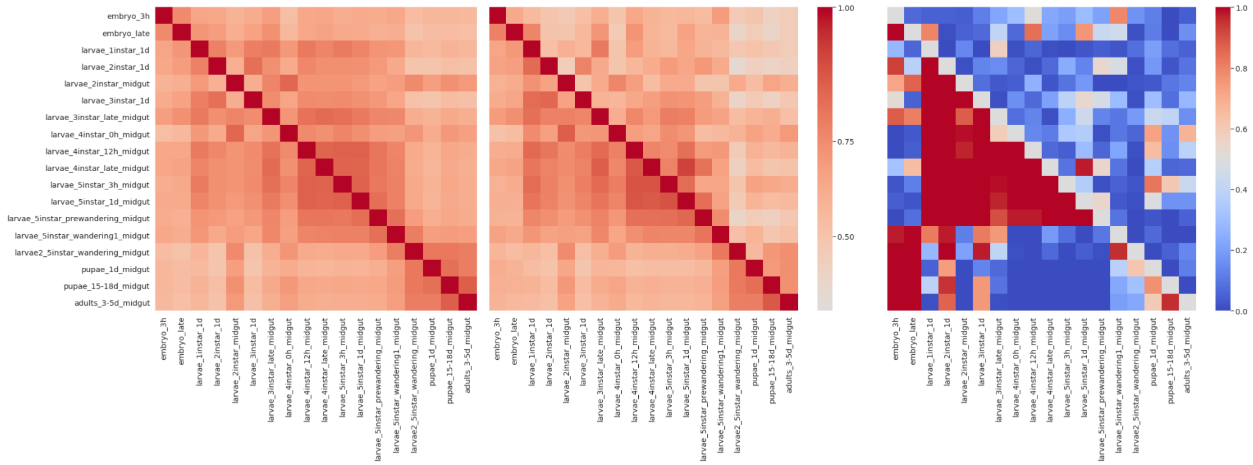

*Manduca sexta*, fat body samples

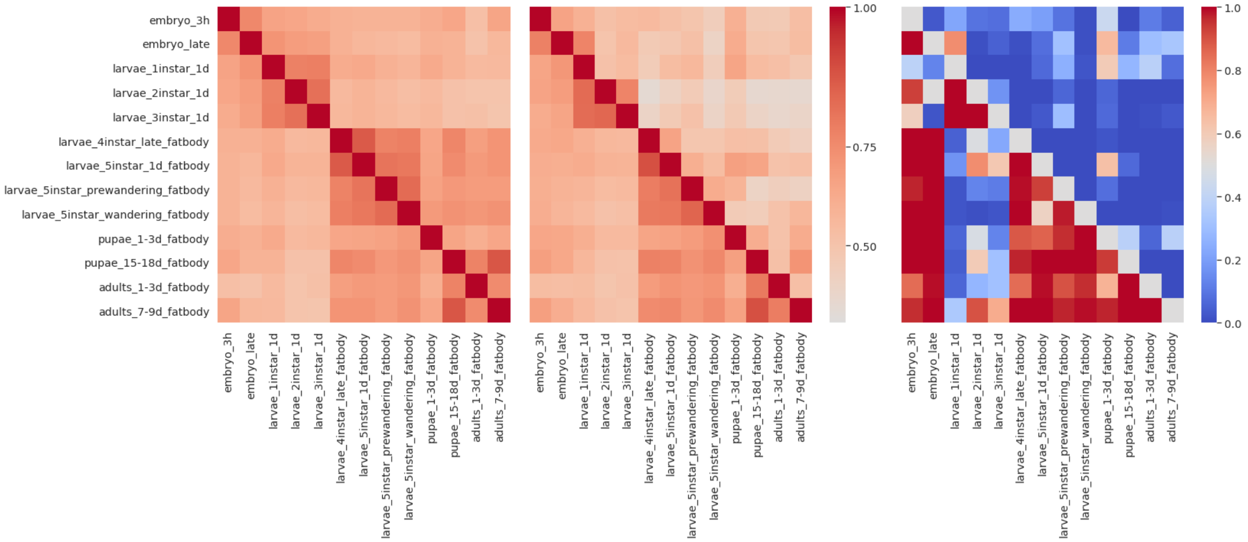


*Polypedilum vanderplanki*

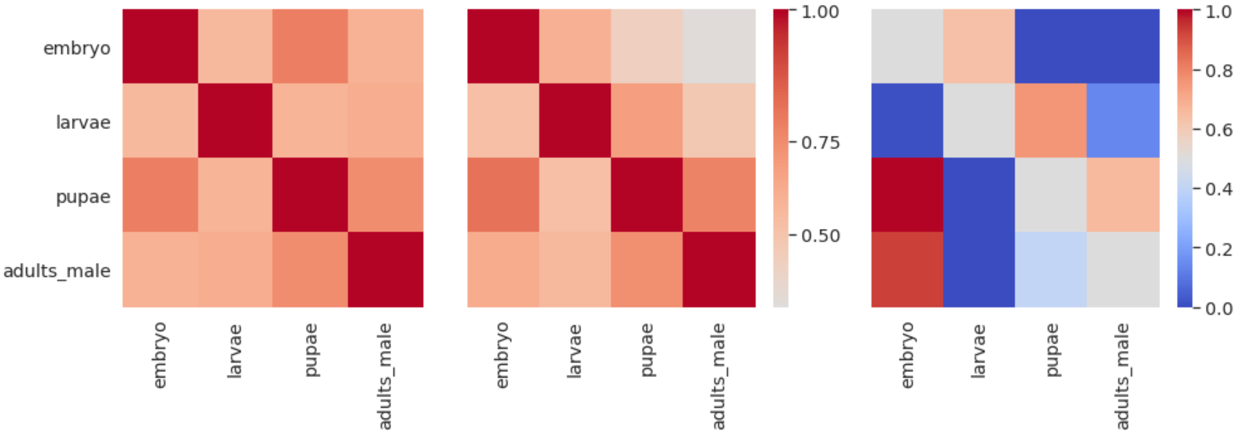
