## Supplementary Figure 2 for "Recapitulation of the embryonic transcriptional program in insect pupae"

### Supplementary Figure 2. Gene expression patterns across the development.

Genes from datasets with more than four measured time points were hierarchically clustered with the Spearman correlation coefficient as the distance metric.

*Drosophila melanogaster*, Daines

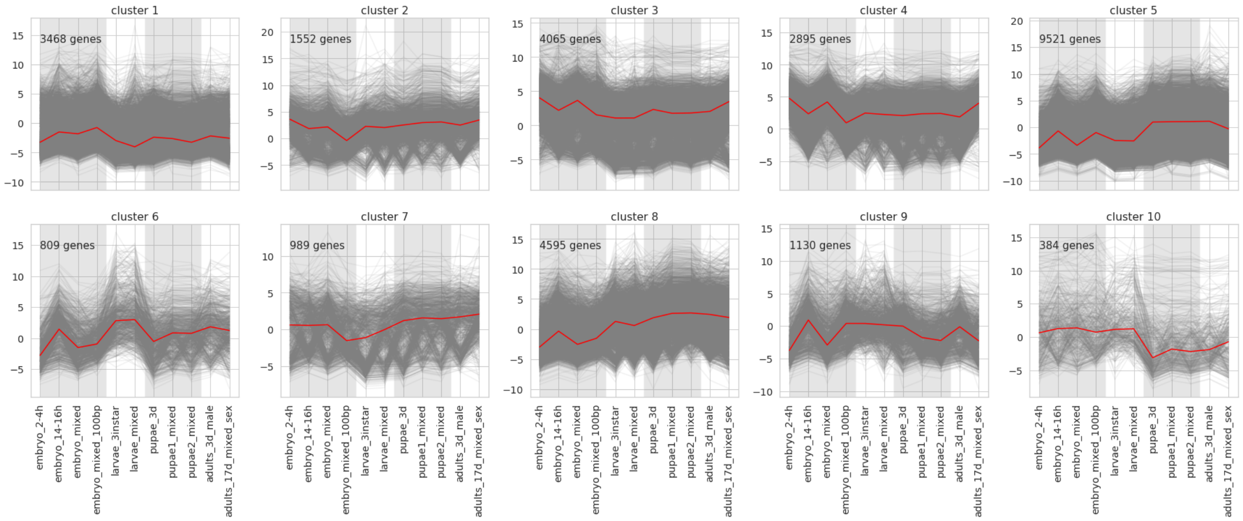


*Drosophila melanogaster*, Graveley

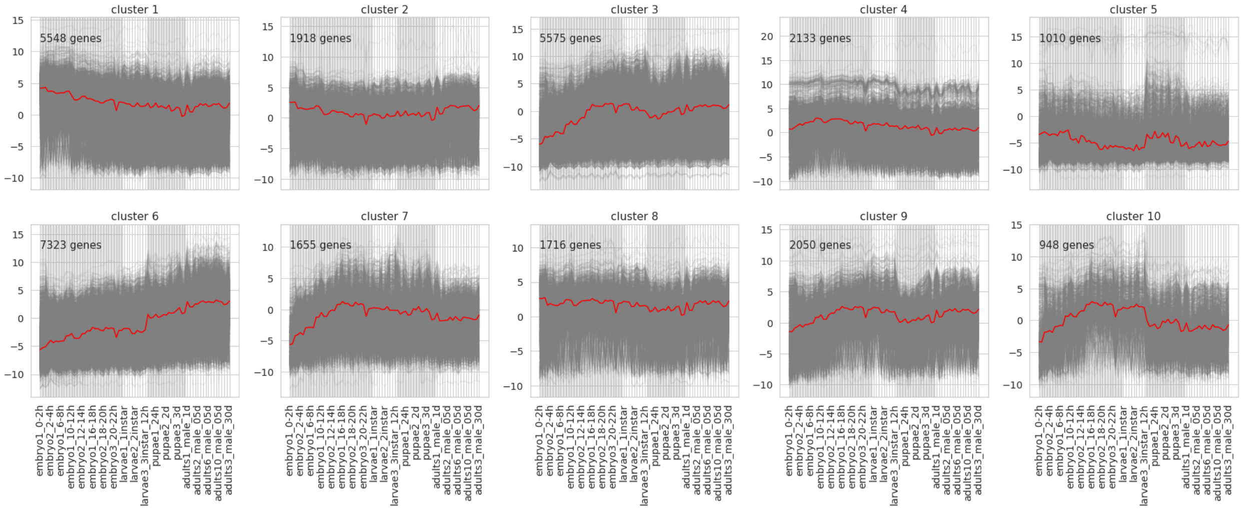

#### *Drosophila melanogaster*, Arbeitman, GPL2837

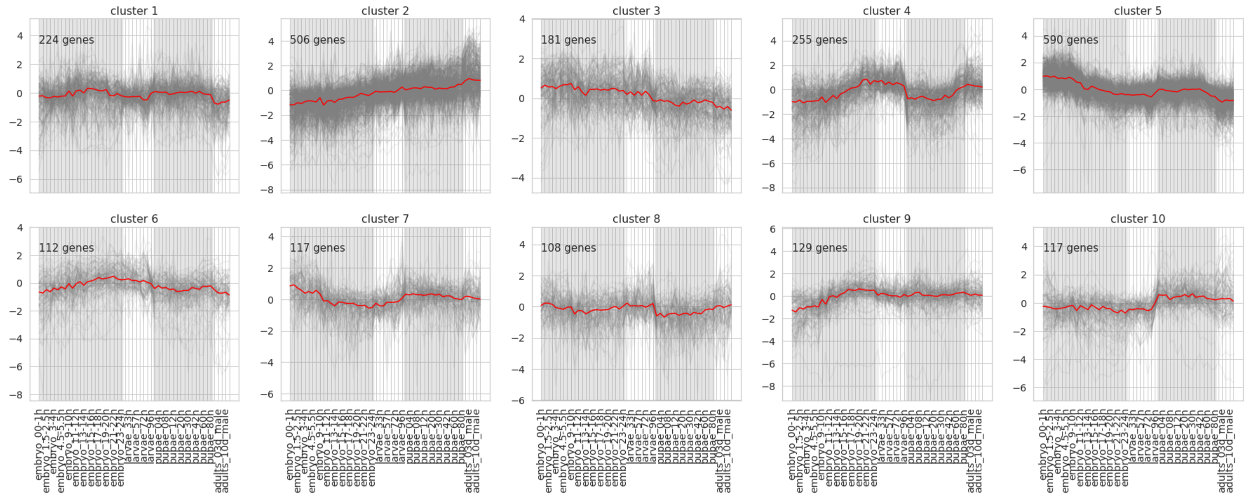


#### *Drosophila melanogaster*, Arbeitman, GPL2838

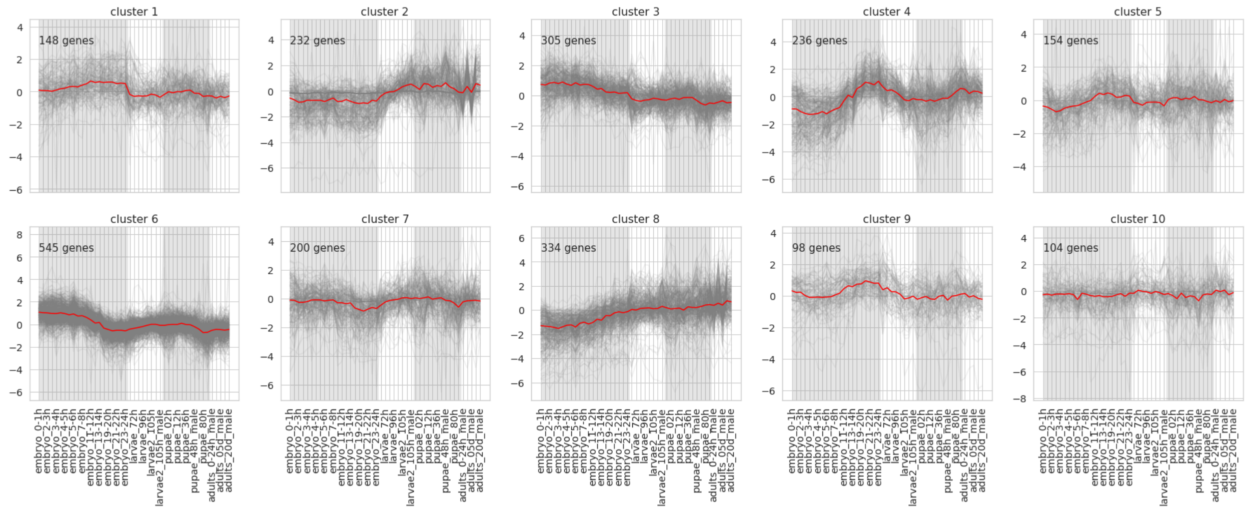


#### *Drosophila melanogaster*, Arbeitman, GPL2840

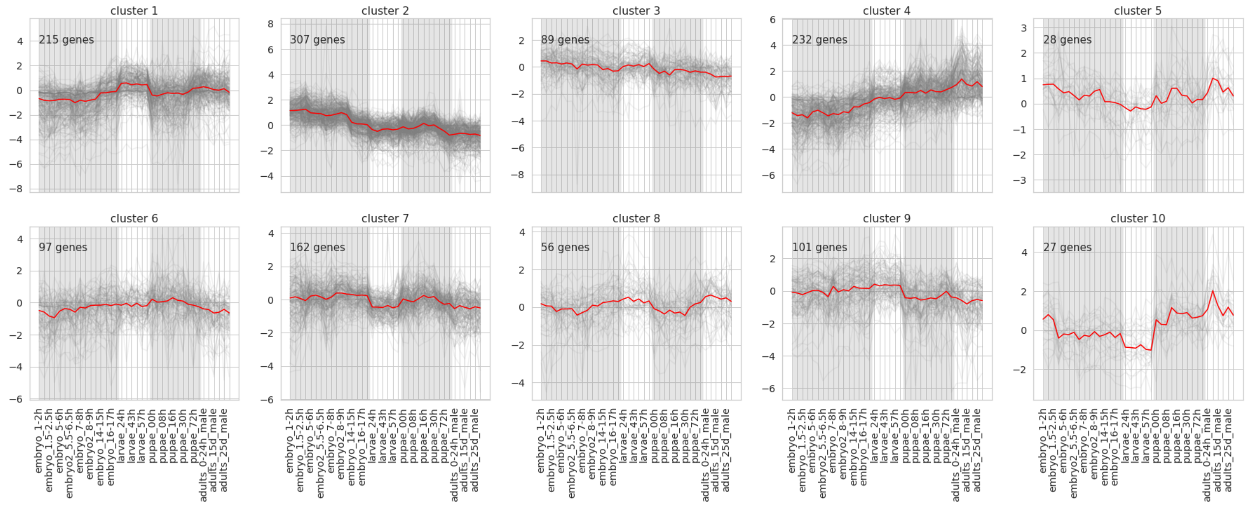

*Zeugodacus cucurbitae*, Wei

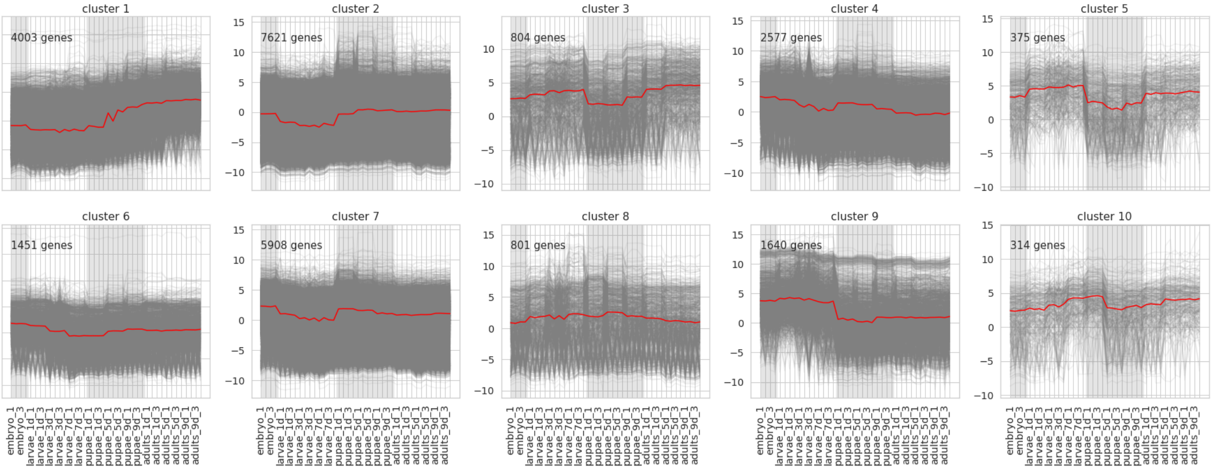


*Manduca sexta*, midgut samples

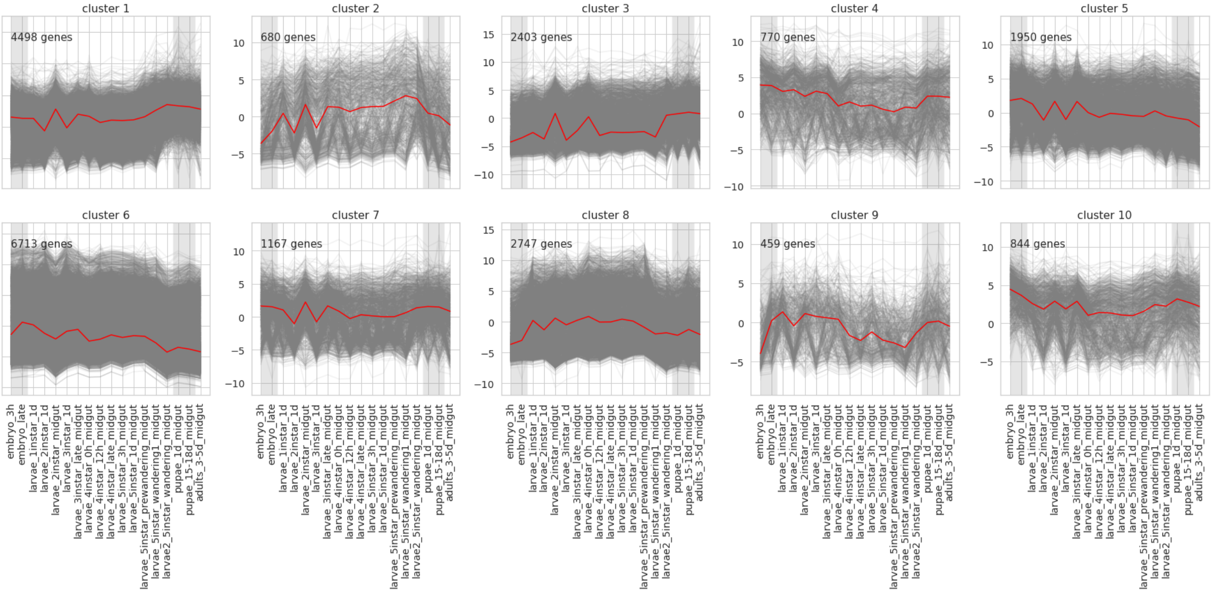

### *Manduca sexta*, fat body samples

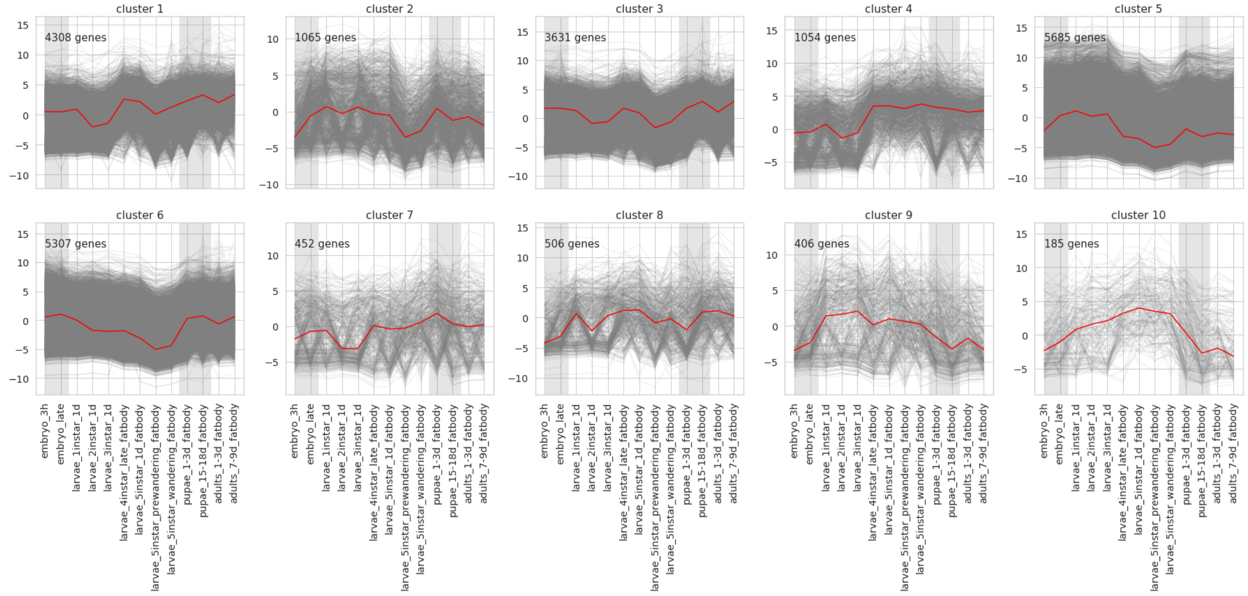
